# Hybrid immunity shifts the Fc-effector quality of SARS-CoV-2 mRNA vaccine-induced immunity

**DOI:** 10.1101/2022.06.10.495727

**Authors:** Kathryn A. Bowman, Daniel Stein, Sally Shin, Kathie G. Ferbas, Nicole H Tobin, Colin Mann, Stephanie Fischinger, Erica Ollmann Saphire, Douglas Lauffenburger, Anne W. Rimoin, Grace Aldrovandi, Galit Alter

## Abstract

Despite the robust immunogenicity of SARS-CoV-2 mRNA vaccines, emerging data reveal enhanced neutralizing antibody and T cell cross-reactivity among individuals that previously experienced COVID-19, pointing to a hybrid immune advantage with infection-associated immune priming. Beyond neutralizing antibodies and T cell immunity, mounting data point to a potential role for additional antibody effector functions, including opsinophagocytic activity, in the resolution of symptomatic COVID-19. Whether hybrid immunity modifies the Fc-effector profile of the mRNA vaccine-induced immune response remains incompletely understood. Thus, here we profiled the SARS-CoV-2 specific humoral immune response in a group of individuals with and without prior COVID-19. As expected, hybrid Spike-specific antibody titers were enhanced following the primary dose of the mRNA vaccine, but were similar to those achieved by naïve vaccinees after the second mRNA vaccine dose. Conversely, Spike-specific vaccine-induced Fc-receptor binding antibody levels were higher after the primary immunization in individuals with prior COVID-19, and remained higher following the second dose compared to naïve individuals, suggestive of a selective improvement in the quality, rather than the quantity, of the hybrid humoral immune response. Thus, while the magnitude of antibody titers alone may suggest that any two antigen exposures – either hybrid immunity or two doses of vaccine alone - represent a comparable prime/boost immunologic education, we find that hybrid immunity offers a qualitatively improved antibody response able to better leverage Fc effector functions against conserved regions of the virus.

## Introduction

Despite the development of several highly protective COVID-19 vaccines, SARS-CoV-2 continues to spread across the globe due to incomplete global distribution of vaccines, waning immunity, as well as the evolution of variants of concern ^1,2^. Currently, only 64.8% of the global population has received at least one dose of vaccine ^3^ and strategic boosting has been complicated by our incomplete understanding of the correlates of immunity against COVID-19 ^4,5^. While neutralizing antibodies clearly contribute to the blockade of viral transmission ^6^, persistent vaccine-induced protection against severe disease and death from several neutralization resistant variants of concern supports the critical role for alternate vaccine induced immunologic responses as key determinants of protection against disease. While T cells have been proposed in the control and clearance of infection after transmission, their direct association with disease severity remains unclear. Conversely, antibodies able to leverage the antiviral function of the immune response, via Fc-receptors, were associated with attenuated symptomatology ^7^, survival of severe COVID-19 ^8^, were conserved for long periods of time ^9^, and maintained function across variants of concern (VOCs) ^10^. Non-neutralizing Fc-effector functions are important in protection against Influenza virus ^11,12^, Ebola virus ^13^, as well as several bacterial infections ^14,15^. These data support a critical role of these alternative antiviral functions of the humoral immune response to SARS-CoV2.

Real world vaccine efficacy revealed rapidly waning immunity following vaccination ^16–18^ prompting recommendations for booster vaccine doses 4 to 6 months following the primary vaccine series ^19,20^. However, anecdotal studies suggested fewer vaccine breakthroughs ^21–24^ and a slower decay in the antibody response ^25^ among individuals who had previously experienced COVID-19 prior to vaccination. Moreover, deeper immunological profiling pointed to increased breadth and magnitude of the neutralizing antibody response in individuals with hybrid (infection + vaccination) compared to vaccine-only induced immunity^26–31^. Similarly, individuals with hybrid immunity (infection + vaccination) produced a distinct population of functionally Th1-skewed IFNγ and IL-10-expressing memory CD4+ T cells ^32^ and CD8+ T cells ^33^ not observed in previously naïve individuals. However, whether hybrid immunity also enhanced the Fc-effector profile of the vaccine induced SARS-CoV-2 specific humoral response remained largely unknown.

As worldwide vaccination efforts continue, a much larger percentage will have previously recovered from natural infection prior to completing vaccination. Thus, understanding the impact of hybrid immunity on shaping the overall humoral immune response may provide key insights on correlates of immunity and guide boosting recommendations. Thus, here we comprehensively profiled the Fc landscape of mRNA induced humoral immune responses across a cohort of individuals that had previously experienced COVID-19 or were infection-naive.

Consistent with prior observations ^26,28,34^, we saw that SARS-CoV-2 vaccine specific titers increased in both the hybrid immunity and infection naïve groups after initial vaccine dose, albeit with higher titers in the hybrid immunity group. As seen in these prior studies ^28,31^, previously infected individuals developed vaccine-induced responses after a single dose of either Pfizer BNT162b2 or Moderna mRNA-1273 mRNA vaccine that were similar in magnitude to antibody responses after two vaccine doses in infection-naïve individuals. Conversely, we observed a significant increase in Fc-receptor (FcR) binding in previously infected individuals after the first dose, that was further expanded after the second dose, potentially conferring broader functional protection against future infection. Thus, hybrid immunity may confer a gain in quality rather than quantity of the antibody response, not apparent on evaluation of titers or neutralizing capacity alone.

## Methods

### Study population

Health services workers at the University of California, Los Angeles (UCLA) and first responders in the Los Angeles County Fire Department (LACoFD) were enrolled in a longitudinal cohort study assessing rates of SARS-CoV-2 infection in high-risk individuals (Table 1) ^35^. Eligible participants were over 18 years of age and free of symptoms associated with COVID-19 prior to enrollment. Participants were asked to provide monthly blood samples and up to biweekly self-collected mid-turbinate nasal swabs. Enrollment began in April 2020. In December 2020, two companies, Pfizer-BioNTech and Moderna, were granted emergency use authorizations (EUA) in the United States of America for their mRNA-based SARS-CoV-2 vaccines encoding the spike (S) protein. Both UCLA Health and LACoFD began offering vaccines in December 2020. Participants had blood drawn between 7 days after the first vaccine dose and just prior to the second dose (up to 20 days after the first dose of BNT162b2 and up to 27 days after the first dose of mRNA-1273). Blood was also collected 7-30 days, 31-60 days and 61-90 days after completion of the two-dose series. All samples were collected between June 29, 2020 and March 11, 2021. Not all participants provided blood samples at every time point. Only individuals receiving the Moderna or Pfizer-BioNTech vaccine and who had been infected prior to the first dose or not at all were retained for analysis (n=63). In particular, four individuals who had become infected between doses were excluded.

**Table 1.**
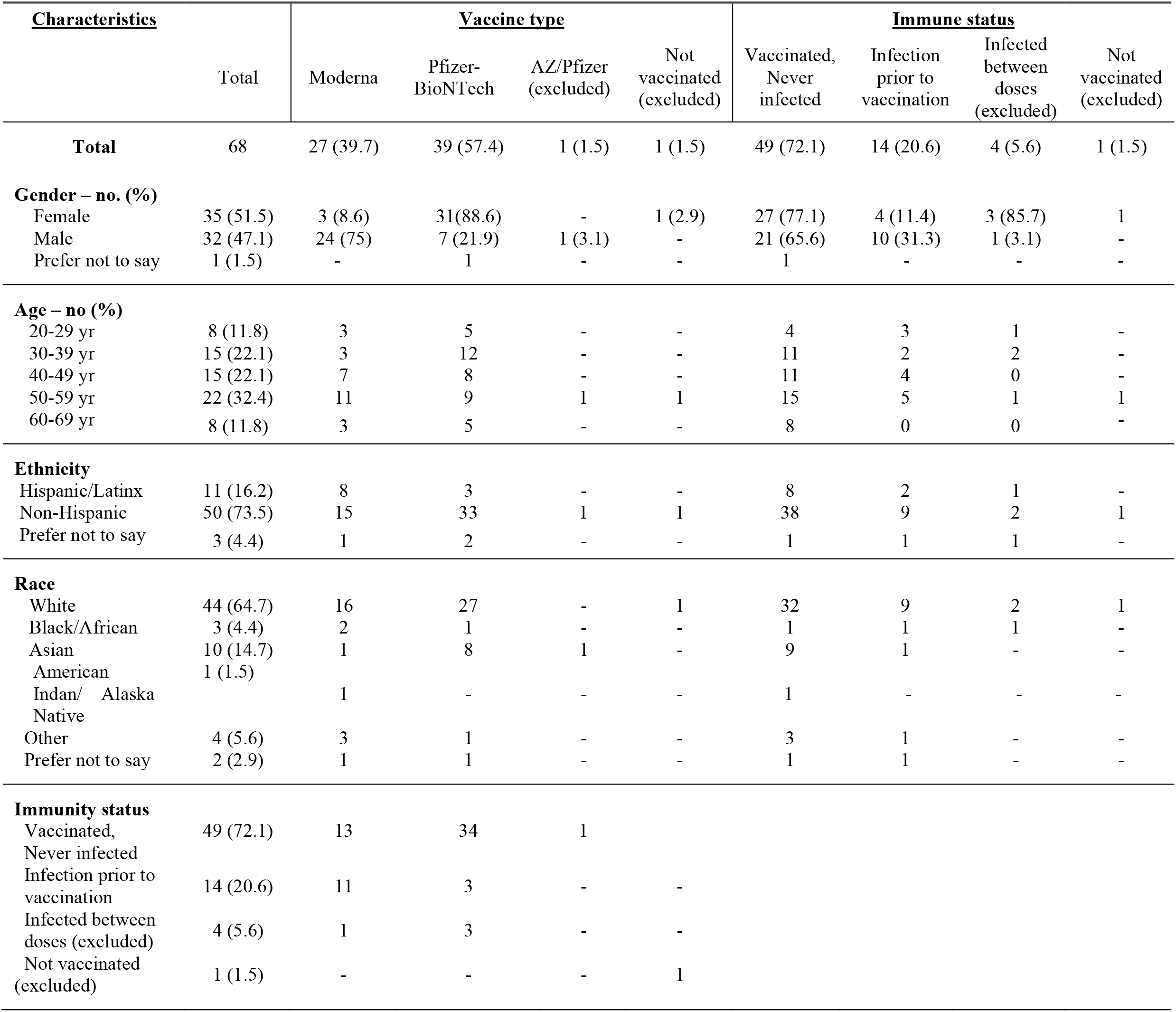
Baseline characteristics and demographic data of the study population.

### Antigens

Antigens used for Luminex based assays: SARS-CoV-2 D614G WT S (provided by Erica Saphire, La Jolla Institute for Immunology), SARS-CoV-2 S1 (Sino Biological), SARS-CoV-2 S2 (Sino Biological) and SARS-CoV-2 RBD (kindly provided by Aaron Schmidt, Ragon Institute), and SARS-CoV-2 NTD (Sino Biological).

### Biophysical antibody profiling (Luminex)

Serum samples were analyzed by customized Luminex assay to quantify the relative concentration of antigen-specific antibody isotypes, subclasses, and Fcγ-receptor (FcγR) binding profiles, as previously described ^36,37^. SARS-CoV-2 antigens coupled to Luminex beads with different fluorescences were used to profile SARS-CoV-2 antigen-specific humoral immune responses in a high-throughput, multiplexed assay. Antigens were coupled to magnetic Luminex beads (Luminex Corp) by carbodiimide-NHS ester-coupling (Thermo Fisher). Antigen-coupled microspheres were washed and incubated with plasma samples at an appropriate sample dilution, based on performed dilution curves (1:500 for IgG1, 1:1000 for all low affinity Fcγ-receptors, and 1:100 for all other readouts) for 2 hours at 37°C in 384-well plates (Greiner Bio-One). Unbound antibodies were washed away, and antigen-bound antibodies were detected by using a PE-coupled detection antibody for each subclass and isotype (IgG1, IgG2, IgG3, IgG4, IgA1, and IgM; Southern Biotech), and Fcγ-receptors were fluorescently labeled with PE before addition to immune complexes (FcγR2a, FcγR2b, FcγR3a, FcγR3b; Duke Protein Production facility). Plates were incubated for one hour, washed, then Flow cytometry was performed on an iQue (Intellicyt), and analysis was performed on IntelliCyt ForeCyt (v8.1). Relative antigen-specific antibody titers are reported as PE Median Fluorescent Intensity (MFI).

### Antibody-dependent complement deposition (ADCD)

Antibody-dependent complement deposition (ADCD) was conducted as previously described ^38^. Briefly, SARS-CoV-2 antigens were coupled to magnetic Luminex beads (Luminex Corp) by carbodiimide-NHS ester-coupling (Thermo Fisher). Coupled beads were incubated for 2 hours at 37°C with serum samples (1:10 dilution) to form immune complexes and then washed to remove unbound immunoglobulins. Lyophilized guinea pig complement (Cedarlane) was diluted in gelatin veronal buffer with calcium and magnesium (GBV++) (Boston BioProducts) and added to immune complexes to measure antibody-dependent deposition of C3. C3 was then detected with an anti-C3 fluorescein-conjugated goat IgG fraction detection antibody (Mpbio). Flow cytometry was performed with iQue (Intellicyt) and an S-Lab robot (PAA). ADCD was reported as the median fluorescence of C3 deposition.

### Antibody-dependent cellular (ADCP) and neutrophil (ADNP) phagocytosis

Antibody-dependent cellular phagocytosis (ADCP) and antibody-dependent neutrophil phagocytosis (ADNP) assays were conducted according to previously described protocols ^39,40^. SARS-CoV-2 antigens were biotinylated using EDC (Thermo Fisher) and Sulfo-NHS-LCLC biotin (Thermo Fisher) and coupled to yellow-green (505/515) fluorescent Neutravidin-conjugated beads (Thermo Fisher). Immune complexes were formed by incubating antigen-coupled beads for 2 hours at 37°C with 1:100 diluted serum samples, then washing to remove unbound immunoglobulins. For ADCP, the immune complexes were incubated for 16–18 hours with THP-1 cells (1.25×b10^5^ THP-1 cells/mL) and for ADNP for 1 hour with RBC-lysed whole blood. After incubation, cells were fixed with 4% PFA. For ADNP, RBC-lysed whole blood was washed, stained for CD66b+ (Biolegend) to identify neutrophils, and fixed in 4% PFA. An iQue (Intellicyt) was used to perform flow cytometry to identify the percentage of cells that had phagocytosed beads and the number of beads that had been phagocytosed. Phagocytic function is reported as a Phagocytosis Score (phagocytosis score = % positive cells × Median Fluorescent Intensity of positive cells/10000). Analysis was performed on IntelliCyt ForeCyt (v8.1).

### Batch Correction

Luminex experiments across samples were performed as described previously^41^, linear mixed-effects modeling was used to remove batch effects from the data while preserving the biological sources of variation that are of interest. In particular, for each feature measured, a linear mixed-effects model was fit to the data with terms accounting for fixed group effects *g*_*i*_, random batch effects *b*_*j*_, and random group-specific batch effects (*gb*)_*ij*_:

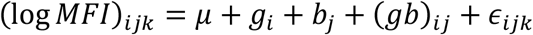

with terms described in Table S1. Linear mixed-effect models were fit using the lme4 package (version 1.1-27.1) in R version 4.1.0^42^. Batch-corrected values were then computed by subtracting the batch effects from each measurement:

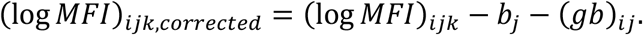

Furthermore, all subsequent analysis on batch-corrected data was confirmed to be consistent with analysis performed on a single batch.

**Table S1:** Terms used in linear mixed-effects model for batch correction

| Term | Variable | Effect Type | Number of Terms |
| --- | --- | --- | --- |
| $(\log MFI)_{ijk}$ | Sample measurement for a single feature | | |
| $\mu$ | Mean log(MFI) for that feature | Fixed | 1 |
| $g_i$ | Group effect of interest<br>(dose and infection status) | Fixed | 4 groups<br>(V1/V2 x infected/not) |
| $b_j$ | Batch main effect | Random,<br>$b \sim N(0, \sigma_b^2)$ | 2 batches |
| $(gb)_{ij}$ | Group x batch interaction | Random,<br>$(gb) \sim N(0, \sigma_{gb}^2)$ | 4 groups x 2 batches = 8 |
| $\epsilon_{ijk}$ | Residual error | Random,<br>$\epsilon \sim N(0, \sigma^2)$ | |

### Univariate Analysis

Univariate statistical analyses were performed using GraphPad Prism version 9.1.2 for macOS. For each feature, comparisons were made between the four vaccinee groups — previously infected and naïve individuals after their first or second doses — using a non-parametric Kruskal-Wallis test followed by Dunn’s multiple comparisons test between all pairs of groups.

### Multivariate Analysis

Multivariate analysis was performed in R (version 4.1.0). Measurements for each feature were first log10-transformed and then centered and scaled. For classification models, LASSO feature selection was performed using the systemsseRology R package (version 1.1) (https://github.com/LoosC/systemsseRology) ^43^. LASSO selection was repeated 50 times, and features that were selected in at least 20% of trials were kept. To discriminate antibody features from previously infected and naïve individuals, partial least squares discriminant analysis (PLSDA) was performed using the LASSO-selected features. The importance of individual features was assessed using variable importance in projection (VIP) scores^44^. Model performance and robustness were assessed using cross-validation, which was compared to control models trained with random features or permuted labels. Separate models were built for each timepoint to distinguish samples from previously infected and naïve individuals, and recipients of Moderna and Pfizer-BioNTech vaccines were combined in the analysis. A LASSO-PLSDA model was also built to distinguish antibody features at different timepoints using all the samples.

For correlation analyses, Spearman correlations were computed between all pairs of antibody features, and multiple comparisons were handled using a Benjamini-Hochberg correction with FDR < 0.05 ^45^. In order to compare correlations between samples from previously infected and naïve individuals, a Fisher’s Z-transformation of the correlation coefficients was computed ^46^. Testing for significance was performed using bootstrap simulations with the bootcorci R package (version 0.0.0.9000) (https://github.com/GRousselet/bootcorci) ^47^.

## Results

### Vaccination-induced antibody response in previously infected and naïve individuals

We comprehensively profiled the SARS-CoV-2 humoral immune response in a group of mRNA vaccinees, including 14 previously infected individuals with SARS-CoV-2 and 49 that were naïve to SARS-CoV-2. The group included health care workers between the ages of 26-68, 35:32 women:men, 39 individuals received the Pfizer/BNT16b2 vaccine and 27 individuals received the Moderna mRNA-1273 vaccine. Vaccine responses were profiled both after the first dose of vaccination (prime, V1) and/or after the second dose (boost, V2) to compare the magnitude, quality, and kinetics of humoral responses after prime and boost in hybrid immunity and infection-naïve individuals. Antibody subclass, isotype, Fc-receptor binding levels, and antibody effector responses were profiled across the full D614G Spike, the S1-domain, the receptor binding domain (RBD), the S2 domain, and the N-terminal domain (NTD). Additionally, the antibody Fc region was simultaneously profiled by measuring isotype-specific responses (IgG1, IgG2, IgG3, IgG4, IgM, IgA1) and FcγR binding (FcγR2a, FcγR2b, FcγR3a, FcγR3b).

Subsequently, Spike-specific antibody effector functions including antibody dependent cellular phagocytosis (ADCP), antibody dependent neutrophil phagocytosis (ADNP), and antibody dependent complement deposition (ADCD) were profiled (**Figure 1A**).

**Figure 1:**
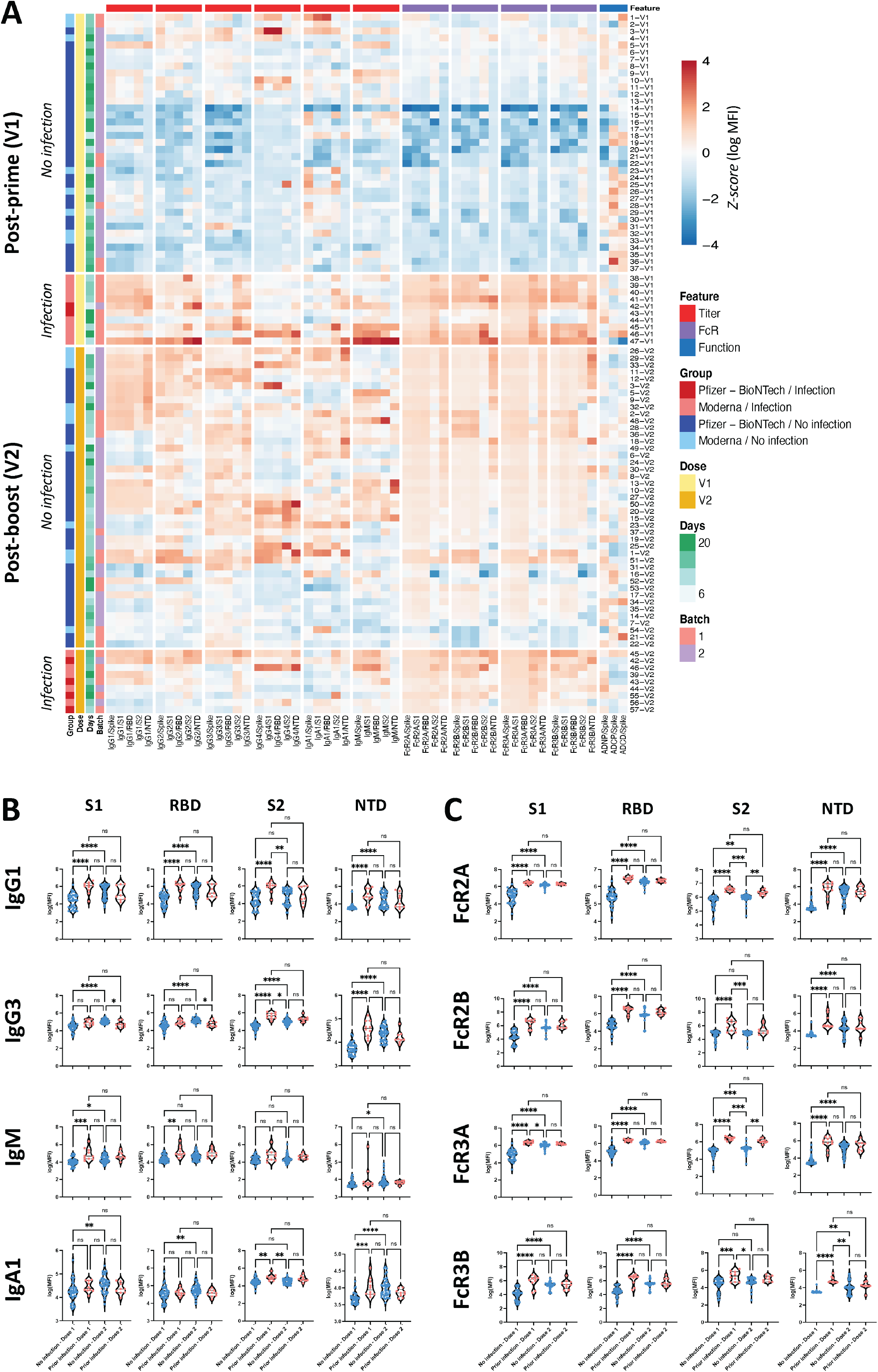
Isotype titers and FcR binding for previously infected individuals after dose 1 are comparable to levels in naïve individuals after dose 2 A. The heatmap shows Z-scored SARS-CoV-2 specific antibody features for individuals post-prime and post-boost. Each row is an individual sample, and each column is a measured feature. Features are clustered hierarchically, and individuals are also clustered within each group (infected / not infected and Pfizer-BioNTech / Moderna). Z-scores are calculated across all samples and truncated between -4 and 4. B. Violin plots show individual subclasses specific for S1, RBD, S2, and NTD for naïve infected and previously infected individuals after 1^st^ and 2^nd^ vaccine doses. Significance was determined by a non-parametric Kruskal-Wallis test followed by Dunn’s multiple comparisons test between pairs of groups. * p < 0.05, ** p < 0.01, *** p < 0.001, **** p < 0.0001. C. Violin plots show low affinity FcR binding by S1, RBD, S2, and NTD-specific antibodies for naïve and previously infected individuals after 1^st^ and 2^nd^ vaccine doses. Significance was determined by a non-parametric Kruskal-Wallis test followed by Dunn’s multiple comparisons test between pairs of groups. * p < 0.05, ** p < 0.01, *** p < 0.001, **** p < 0.0001.

Consistent with prior observations, after the first immunization, individuals with prior SARS-CoV-2 infection induced higher levels of S1-, RBD- and S2-specific IgM, IgG1, and IgG2 compared to naïve individuals (**Figure 1B**). Conversely, there was minimal difference in the IgG3 and IgA response across the groups, and limited to S2-specific responses. Interestingly, a single dose of mRNA did not induce appreciable NTD-specific antibodies across the naïve vaccinees, but did induce robust NTD-specific IgG1, IgG2, IgG3, and IgA in individuals that were previously infected (**Figure 1B**). After the second dose of mRNA, S1-specific IgG1, IgG2, IgG3, IgM, and IgA antibody titers rose significantly in vaccinated individuals, reaching levels equal or superior to those found in previously infected individuals. Unexpectedly, NTD- and S2-specific responses exhibited a declining trend among previously infected individuals after the second dose. Thus, overall, the broader epitope-specific recognition observed in previously infected individuals diminished with a boost, with similar isotype titers across the two groups after boosting.

Beyond binding, antibody interactions with Fc-receptors (FcRs) drive antibody effector functions ^48^. Thus, to understand whether prior infection shapes the functional potential of the mRNA vaccine-induced SARS-CoV-2 specific humoral immune response, we probed the ability of vaccine-induced antibodies to interact with the 4 low affinity Fc-receptors in humans (FcγR2a, FcγR2b, FcγR3a, and FcγR3b) that largely direct innate immune effector functions ^48,49^. Across all antigens, elevated FcR binding was noted in individuals that had previously been infected after the primary mRNA vaccine dose (**Figure 1C**). After the second dose, naïve vaccinees experienced a significant rise in FcR binding to all FcγRs. Previously infected vaccinees did not experience any further maturation of the functional humoral immune response, although FcR binding remained higher for S2-specific FcγR2a and FcγR2b binding in previously infected individuals than the naïve individuals after second dose. Viewed as a whole, there are clear qualitative differences in the architecture of the hybrid vs the naïve immune response after the first and second dose of vaccine, not fully explained by a net number of antigen exposures.

### Previous infection expands the epitope-specific and functional SARS-CoV-2 response

Given the presence of striking univariate differences both in the overall breadth and FcR binding profiles of vaccine induced immune responses across the previously infected and naïve vaccinees, we next aimed to define the features that differed most across the SARS-CoV-2 specific immune profiles (**Figure 2**). Specifically, we aimed to SARS-CoV-2-specific antibody profile differences after the first (V1) and second (V2) dose across the groups. A least absolute shrinkage and selection operator (LASSO) was first used to reduce the features to a minimal set of features that differed across the 2 groups, followed by a partial least squares discriminant analysis (PLSDA) for visualization. Vaccine-induced antibody responses after the first vaccine dose (V1) could be clearly separated between previously infected and naïve individuals (**Figure 2A**, p < 0.001, permutation test; LOO-CV accuracy = 1.0) based on as few as 4 of the total 53 features collected per plasma sample (**Figure 2B**). The 4 features were enriched among previously infected individuals and included S2-specific FcγR3a, RBD-specific FcγR3a, S2-specific IgG3, and NTD-specific IgG1 responses, pointing to increased breadth of binding across the S2 and NTD as well as enhanced FcγR3a binding activity among previously infected individuals.

**Figure 2.**
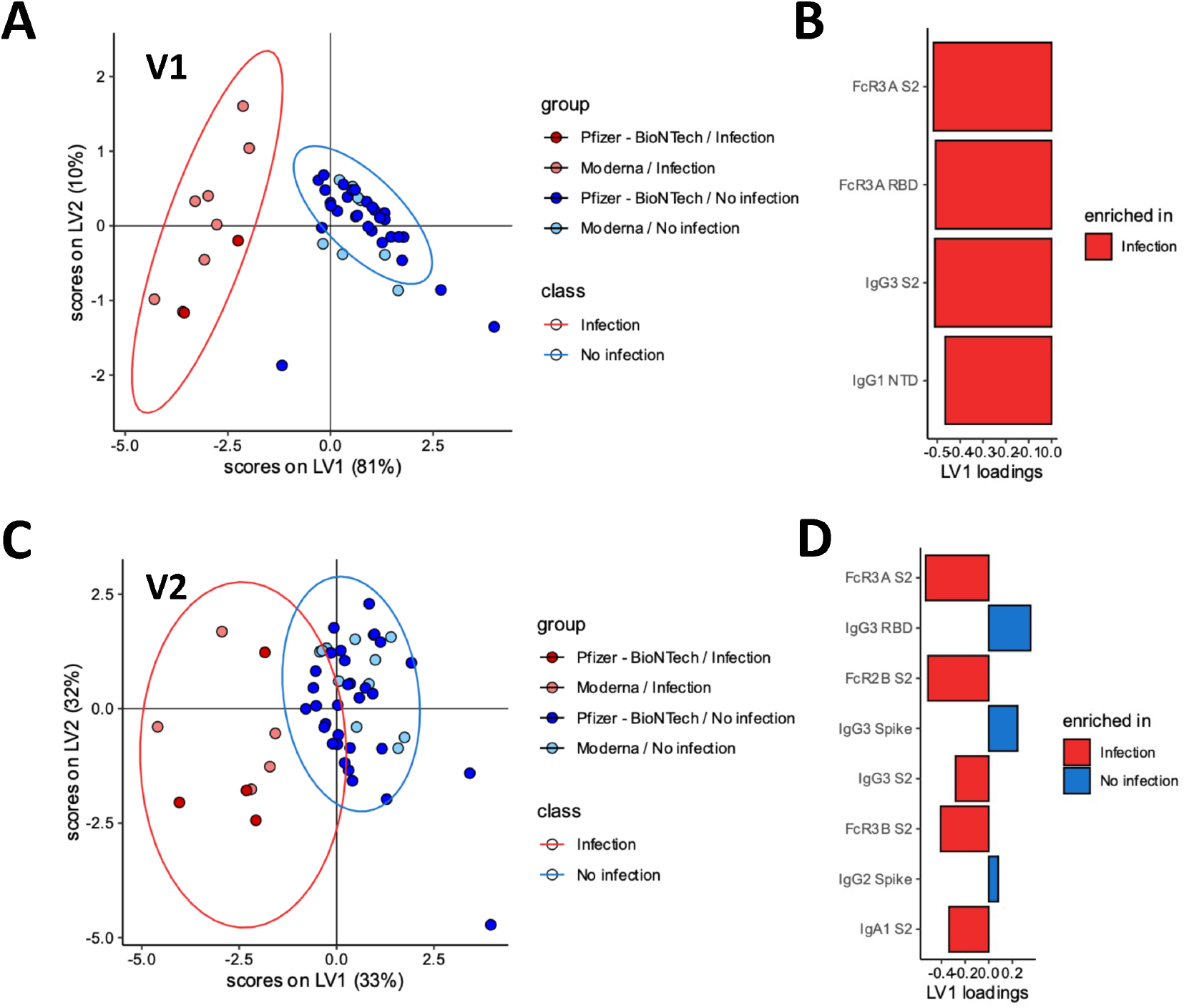
Differences in antibody response following mRNA vaccination for previously infected individuals. A. For the V1 data from all groups, a PLSDA model was constructed to distinguish previously infected individuals from infection-naïve individuals. In the model, individuals receiving the Pfizer-BioNTech and Moderna vaccines were combined within each class (prior infection vs. infection-naive). Scatterplot shows latent variable scores, where each point represents a single individual. Ellipses represent 95% confidence intervals for each class. B. LASSO selection from the PLSDA model of V1 data from all groups. Bar plots show the contribution of each LASSO-selected antibody feature to the latent variables, sorted by VIP score (most important at the top). Bar color corresponds to the class with greater mean value for that feature. C. For the V2 data from all groups, a similar PLSDA model was constructed as in A, to distinguish previously infected individuals from infection-naïve individuals. The scatterplot showls latent variable scores. Ellipses represent 95% confidence intervals for each class. D. LASSO selection from the PLSDA model of V2 data from all groups. Bar plots show the contribution of each LASSO-sselected antibody feature to the latent variables, sorted by VIP score.

After the second dose, previously infected individuals still harbored a distinct SARS-CoV-2 specific antibody profile (**Figure 2C**, p < 0.001, permutation test; LOO-CV accuracy = 0.981) compared to naïve individuals. A total of 8 features were required to achieve separation between the 2 vaccine profiles, marked by 5 features that were elevated in infected individuals and 3 features that were preferentially expanded in naïve individuals (**Figure 2D**). Specifically, S2-specific immunity was overall expanded among the previously infected group of vaccinees, including enhanced titers and FcR binding levels. Conversely, RBD- and Spike-specific IgG3 responses were selectively increased in naïve vaccinees, consistent with the generation of new B cell responses in the setting of a more naïve system, biased to the RBD and S1 domains.

### Differences in coordination between antibody features for previously infected individuals

Given the significant differences in antibody responses across the previously infected and naïve vaccinees, we finally aimed to determine whether previous infection may drive a unique coordination in the humoral immune response induced by the SARS-CoV-2 vaccine (**Figure 3**). As expected, we observed strong positive correlations between SARS-CoV-2-specific IgG titers and FcR-binding among both infected and naïve individuals (**Figure 3A**). After the first vaccine dose (V1), enhanced correlation was observed between IgG3 responses and FcR binding among naïve individuals that was not clearly observed in previously infected individuals (**Figure 3A**). IgM in previously infected individuals was more weakly correlated with FcR binding than in naïve individuals. Moreover, after the first dose, a differential heatmap showed that IgG titers were more robustly correlated with antibody-dependent neutrophil phagocytic (ADNP) activity, whereas FcR-binding levels were more robustly associated with ADNP in naïve individuals (**Figure 3B**).

**Figure 3.**
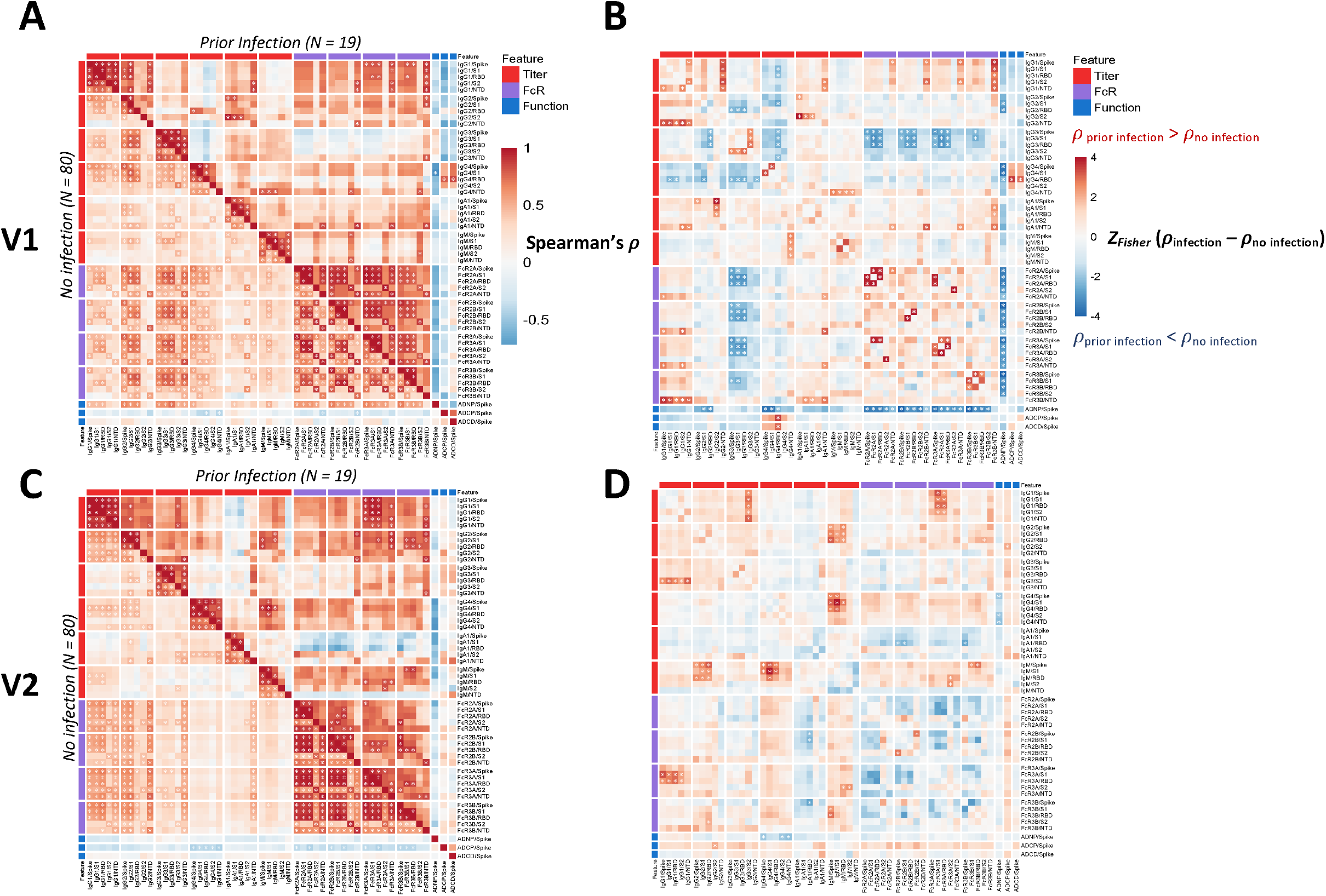
Coordination between antibody isotypes, FcR binding, and antibody functions differs in previously infected individuals A. Spearman rank correlations *ρ* are shown between all pairs of features for previously infected individuals (above diagonal) and for naïve individuals (below diagonal), combining samples from V1. Significant correlations after Benjamini-Hochberg correction with FDR < 0.05 are indicated with *. B. Differences *ρ*_infection_ – *ρ*_no infection_ between correlations among previously infected and among naïve individuals are shown for V1, where blue indicates more positive correlations among naïve individuals. Correlation differences were computed using Fisher’s Z-transformation, and significance was determined using bootstrap simulations. C. Spearman rank correlations *ρ* are shown between all pairs of features for previously infected individuals (above diagonal) and for naïve individuals (below diagonal), combining samples from V2. Significant correlations after Benjamini-Hochberg correction with FDR < 0.05 are indicated with *. D. Differences *ρ*_infection_ – *ρ*_no infection_ between correlations among previously infected and among naïve individuals are shown for V2, where blue indicates more positive correlations among naïve individuals. Correlation differences were computed using Fisher’s Z-transformation, and significance was determined using bootstrap simulations.

After V2, previously infected individuals continued to demonstrate poor correlation between IgG3 and FcR binding, as well as a weaker correlation between IgA1 and FcR binding compared to infection naïve individuals (**Figure 3C**). Conversely, naïve individuals had poor correlation between subclass titers and FcR binding, and notably weaker correlation of S2-specific FcR engagement compared with previously infected individuals. ADNP activity was positively correlated across IgG subclasses and FcRs in naïve individuals and negatively correlated across FcR binding in previously infected individuals, suggesting a shift in previously infected individuals away from ADNP as a dominant effector function. Moreover, significant differences were noted in the correlations between FcR binding in ADNP across the groups, with IgG3-driven ADNP in previously infected individuals and a more diffuse role of FcR-binding antibodies in driving ADNP among naïve vaccinees. Inverse differences were observed in ADCD-driving antibodies, driven by a diffuse set of FcR binding antibodies among previously infected vaccinees, but IgA and IgM in naïve vaccinees with a largely negative correlation of ADCD activity across IgG subclasses and FcγRs. Moreover, the differential heatmap highlighted the presence of an overall shift towards more coordination in IgG titers and FcR binding and function among previously infected individuals, with an overall pink shade in the differential heatmap (**Figure 3D**), marked by enhanced coordination of IgG1 titers with FcγR3a binding, enhanced coordination of antibody features with antibody dependent cellular monocyte phagocytosis (ADCP), NTD-specific IgA levels and antibody dependent complement deposition (ADCD) in previously infected individuals. Conversely, FcR-binding was more highly coordinated across FcRs, across antigen-specificities among the naïve individuals. Together, these results further highlight qualitative differences in antibody responses following vaccination in previously infected individuals.

## Discussion

Due to the slow global rollout of vaccines, much of the world will have experienced SARS-CoV-2 infection prior to immunization ^50^. Along these lines, emerging data suggest qualitatively superior vaccine-induced humoral and cellular immune responses among individuals who were previously infected with SARS-CoV-2 compared with those that receive the vaccine in the absence of prior exposure to the virus ^26–28,31,32^. Yet, beyond neutralizing antibody and cellular immunity, mounting data point to a potential role for Fc-effector functions in the control and clearance of infection. Specifically, opsinophagocytic activity has been linked to survival of severe disease ^51^, enhanced FcR binding has been linked to asymptomatic infection ^8,10^, Fc-effector function has been linked to convalescent plasma therapy ^52^, and Fc-effector function plays a critical role in monoclonal therapeutic activity ^53,54^. However, whether hybrid immunity alters Fc-effector function, potentially resulting in enhanced protection^55^ remained largely unknown.

In our study, we found clear differences in SARS-CoV-2 specific serum antibodies, following vaccination, between previously infected and naïve individuals, with antibody responses of greater magnitude and epitope specificity in individuals vaccinated after prior infection. Although antibody levels approximated each other after V2, several important differences persisted in FcR responses and shifts in coordination of the immune response. We noted that previously-infected individuals had lower RBD-IgG3 levels, pointing to enhanced class-switching, and enhanced FcR binding marking the generation of potentially functionally optimized antibodies particularly targeting the S2-domain of the highly conserved segment of the Spike protein.

Interestingly, enhanced FcR binding in hybrid immunity was particularly skewed towards enhanced binding to FcγR2a and FcγR3a, the two activating FcRs in humans ^56^. Due to its broad expression across immune cell types, particularly on myeloid cells, FcγR2a is poised to drive rapid and robust opsinophagocytosis. Conversely, FcγR3a is expressed in a slightly more restricted manner, on cytotoxic NK cells and mature myeloid cells, implicated in driving rapid cytotoxic granule release and myeloid activity, respectively. Thus, the selective induction and preserved elevated FcγR2a and FcγR3a antibodies may enable individuals with hybrid immunity to clear viruses and kill infected cells more aggressively, providing an advantage even in the face of the emergence of VOCs that evade neutralization.

A specific expansion of S2-specific FcR binding capacity was notable following hybrid immunity. Conversely, vaccination alone drove a S1-dominant response, likely due to the stabilized nature of the Spike antigen in the mRNA vaccines ^57,58^, that likely renders S2 slightly less visible to the immune response. Yet, during viral infection, copious copies of Spike are produced that are presented to the immune system as trimers, monomers, and S1 or S2 components. This heterogeneous presentation likely breaches the stabilized-vaccine immunodominance of S1, providing a unique opportunity for the hybrid immune response to generate immunity to the conserved S2 segment of Spike. Along these lines, the S2 domain is highly conserved across VOCs, with only 6 amino acid substitutions in S2 in Omicron ^59^ as well as high conservation across beta coronaviruses ^60^. Given our emerging appreciation for the disease attenuating, rather than blocking, functions of S2-specific antibodies ^61^ that are mediated largely via Fc-effector functions ^60^, these data argue that hybrid immune induction of potentially cross-reactive, functional antibodies to the S2 may contribute to more robust protection against VOCs.

While this analysis did not capture differences in the quality of the hybrid immune response induced following infection with distinct VOCs, this study highlights an important role for hybrid immunity in driving expanded Fc-effector functions. Thus, beyond improved level of neutralization ^62^ and T cell immunity, induced by hybrid immunity, this study highlights an additional aspect of expanded non-neutralizing antibody Fc-effector function ^28,29^. Future studies evaluating the effects of hybrid immunity on FcR binding and Fc effector functions in the setting of a heterologous combination of Spike exposures, as with an initial VOC infection preceding vaccination, will be critical to understanding the role of potential immune imprinting and the impact of homologous vs heterologous Spike challenge on Fc interactions and downstream effector functions which may play an important ancillary role in protection.

Given the small size of the cohort, we were unable to account for different potential effects of Moderna mRNA-1273 and Pfizer-BNT162b2, nor did we evaluate other non-mRNA vaccine platforms. Despite these limitations, this study highlights valuable qualitative differences in the hybrid immune response not captured in previous studies. While comparable antibody titers after an equal number of total antigenic exposures (natural infection or vaccine) suggest natural infection as an interchangeable priming event with V1 in naïve individuals, this study instead highlights qualitative advantages in the hybrid immune response that may offer potential improvements in vaccine development and merit longer term followup studies to understand the durability of these differences.

## Acknowledgment/Funding

We thank Mark and Lisa Schwartz, Terry and Susan Ragon, and the SAMANA Kay MGH Research Scholars award for their support. Massachusetts Consortium on Pathogen Readiness (MassCPR), the NIH (3R37AI080289-11S1, R01AI146785, U19AI42790-01, U19AI135995-02, U19AI42790-01, 1U01CA260476 – 01, CIVIC75N93019C00052), as well as NIH training grant funding 5T32 AI007387-32.

